# einprot: flexible, easy-to-use, reproducible workflows for statistical analysis of quantitative proteomics data

**DOI:** 10.1101/2023.07.27.550821

**Authors:** Charlotte Soneson, Vytautas Iesmantavicius, Daniel Hess, Michael B Stadler, Jan Seebacher

## Abstract

We describe einprot, an R package providing easy-to-use reproducible workflows for quality control, statistical analysis and visualization of quantitative proteomics data. einprot is applicable to tabular output from MaxQuant, Proteome Discoverer and FragPipe, and a single function call generates an html report that describes the full analysis pipeline applied to the data and contains static and interactive figures and tables for further exploration. This has the potential to facilitate routine analyses as well as to provide a standardized, yet comprehensive way to communicate results to collaborators and the broader community. The source file underlying the report is also returned, giving the user full flexibility to further modify the workflow according to their needs.

## SUMMARY

Quantitative proteomics has come a long way - what used to be specialized analyses performed in proteomics research groups is nowadays a routine service in many proteomics core facilities, and a large collection of sophisticated quantification and analysis tools are available. Yet, the necessary reporting tasks, including statistical analyses of the resulting data, as well as describing all data processing steps, providing quality control, exploration opportunities and result visualizations for publication in a user-friendly way, are generally not routine or automated, and many different analysis workflows are conceivable ^1^. Moreover, additional downstream analyses and integration with other types of data are often necessary, and these are more likely to succeed when all steps of the routine data analysis workflow are transparent and well documented.

The einprot R package provides accessible workflows accommodating quality control, filtering, exploratory analysis and statistical analysis of proteomics data quantified with several commonly used tools, including label-free quantification with MaxQuant ^2^ or FragPipe ^3^, and tandem mass tag (TMT)-multiplexed data quantified with Proteome Discoverer ^4^. Each workflow is provided in the form of a template R Markdown (Rmd) file ^5^, containing the code to be executed as well as text descriptions and explanations of the individual steps. This can significantly reduce the amount of time spent on routine processing tasks, e.g., for a core facility, and enables the entire analysis process to be shared with collaborators or data generators in a way that is comprehensive and easy to follow. In addition, the report contains several interactive tables and plots that allow users to immediately explore their data.

To run a workflow, the user calls a single function in their R session, to which they provide the path to the quantification file(s) as well as a number of additional arguments specifying details about the experiment and the requested analyses. einprot then copies the template Rmd script from the package location to the designated output directory, injects the arguments specified by the user to create a stand-alone file, and compiles this into an html report describing the complete analysis. The stand-alone Rmd file is retained and can, if necessary, be manually modified and fine-tuned by the user and recompiled. Alternatively, the analyst can provide their own template Rmd file if custom analyses are desired. A collection of example reports generated by einprot are provided at https://csoneson.github.io/einprot_examples/. The einprot functions used in the workflows can also be called directly in the R session for an interactive analysis or recreation of specific plots (see the online [vignette](https://fmicompbio.github.io/einprot/articles/einprot.html) for examples). While compiling the report, einprot exports a set of text files and publication-ready plots for further inspection and dissemination of the results (see the [vignette](https://fmicompbio.github.io/einprot/articles/einprot.html) for a full list of output files and Figure 1 and Figure 2 for examples of figures generated by the workflows).

**Figure 1:**
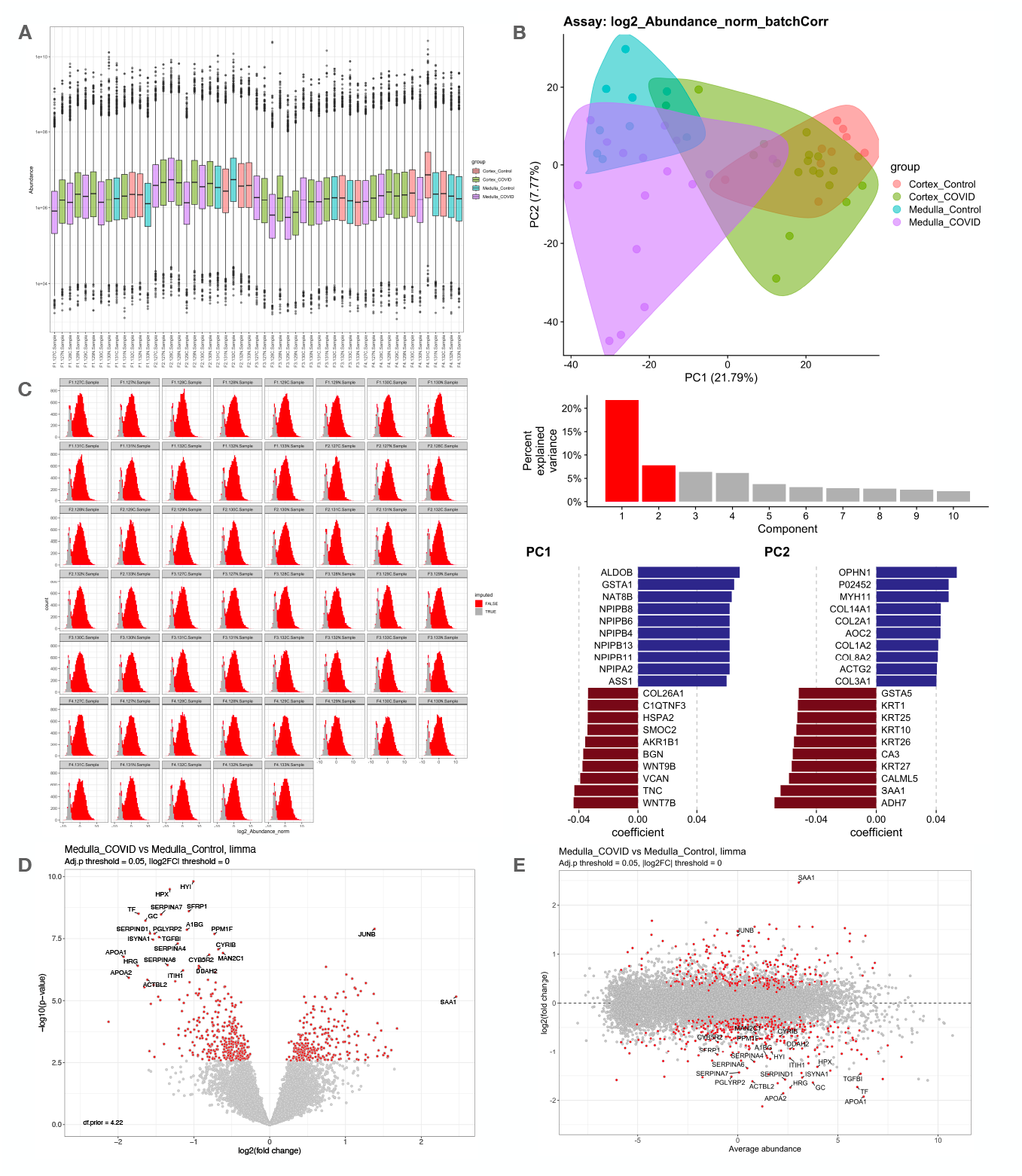
Example figures generated by the einprot workflow, based on a data set from Nie *et al* ^6^, requantified by He *et al* ^7^. The figures can also be generated programmatically from the SingleCellExperiment object generated by the einprot workflow. A. Distribution of abundances in each sample. B. Principal component analysis representation of samples, percent variance explained by each principal component, and the proteins with the highest loadings in the first two components. C. Histograms showing the distribution of observed and imputed abundance values in each sample. D. Volcano plot for one of the tested contrasts. E. MA plot for the contrast shown in D.

**Figure 2:**
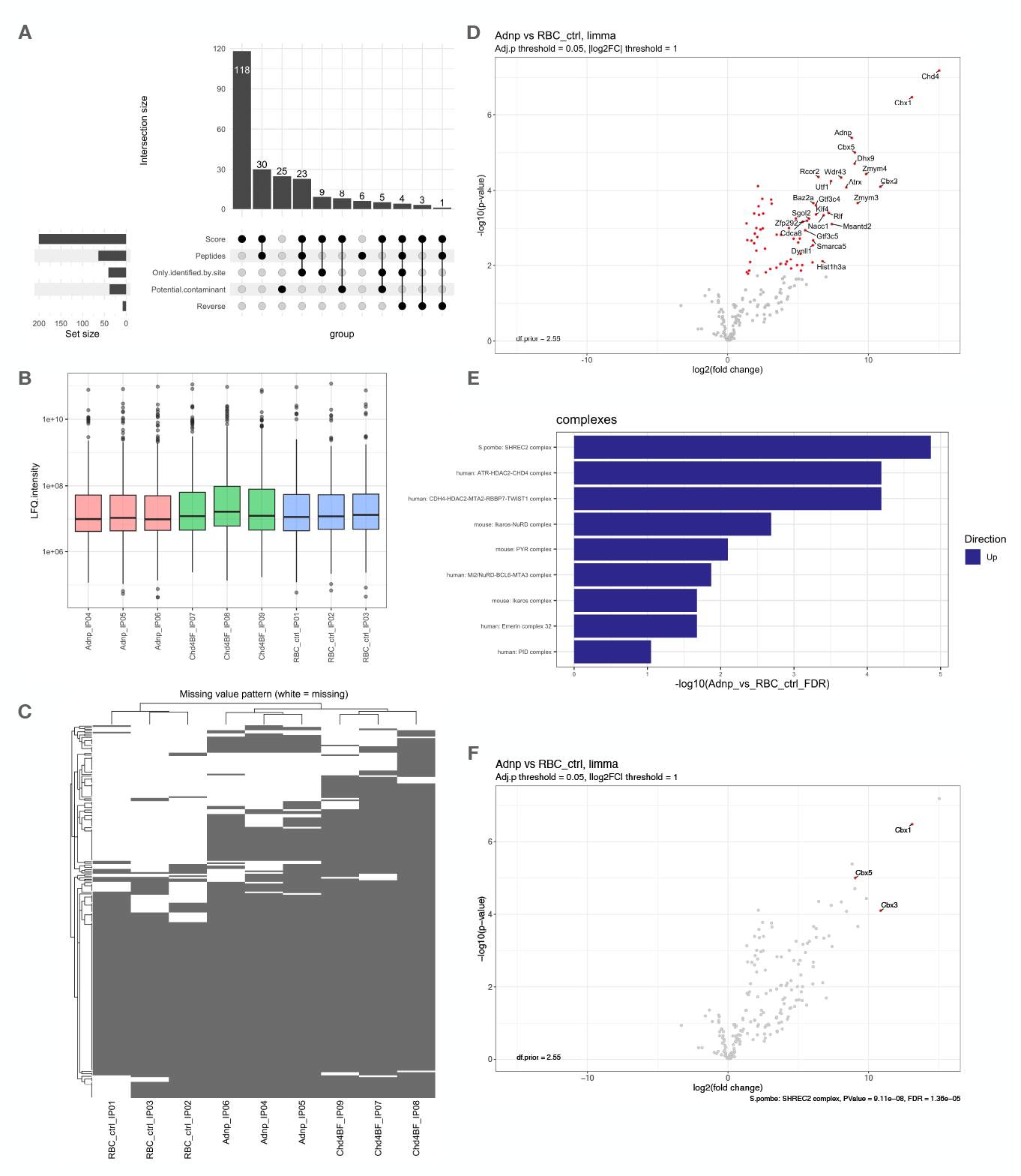
Example figures generated by the einprot workflow, based on an IP-MS data set from Ostapcuk *et al* ^8^, quantified with MaxQuant. A. Overview of the proteins filtered out by the workflow. B. Distribution of LFQ intensities across samples. C. Global missing value pattern. D. Volcano plot for one of the tested contrasts. E. List of known complexes most strongly associated with the contrast in D. F. The same volcano plot as in D, but with the members of the top-ranked complex from E highlighted.

The current version of einprot fully supports five common organisms: *Mus musculus, Homo sapiens, Caenorhabditis elegans, Saccharomyces cerevisiae* and *Schizosaccharomyces pombe*. For these organisms, the user can elect to perform automatic enrichment testing of Gene Ontology terms ^9,10^ and known protein complexes, obtained from CORUM ^11^, PomBase ^12^ and CYC2008^13^ and mapped to the organism of interest using the ortholog mapping from the babelgene R package ^14^. Other species can be analyzed by skipping (parts of) the automatic enrichment testing. Finally, for improved interoperability with other tools, especially from the Bioconductor ecosystem ^15^, einprot stores all raw and processed values (including, e.g., results from differential abundance analysis and dimensionality reduction via principal component analysis) in a SingleCellExperiment object ^16^. In addition, einprot automatically generates an R script that can be sourced to launch a customized interactive exploration session using the iSEE R package ^17^ (Figure 3).

**Figure 3:**
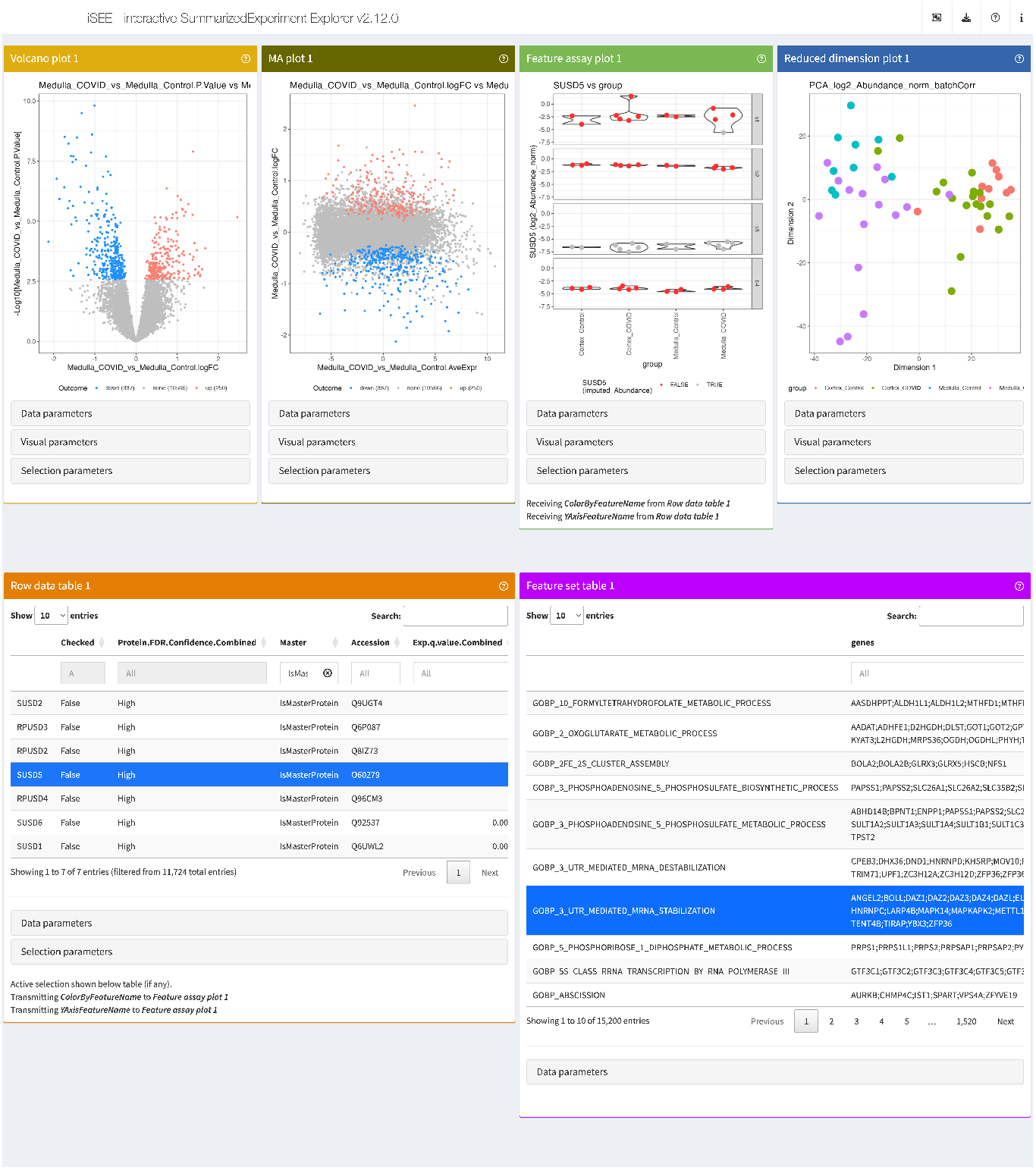
A part of the interactive interface configured for each einprot run, based on the iSEE package. A variety of interactive panels allows the user to explore the statistical results, abundance values and imputation status for individual proteins, a low-dimensional representation of the samples, as well as to browse all results in table form. In addition, all the flexibility provided by the iSEE package is retained, and the user can shape the interface and panel composition according to their own needs.

## STATEMENT OF NEED

Several other toolkits are available for analyzing proteomics data ^18^. Arguably, routine statistical analyses are most commonly performed using vendor-or community-developed software suites accessible via graphical user interfaces, such as Perseus ^19^ or Proteome Discoverer ^4^. While they provide comprehensive analysis capabilities, they are typically not fully open source, and sometimes only available for selected operating systems. In addition, the manual “point-and-click” interface means that they are difficult to incorporate into an automated or programmable data analysis pipeline. Among the broad range of community-developed opensource tools, many (e.g., POMAShiny ^20^, LFQ-Analyst ^21^, amica ^22^, Eatomics ^23^, ProVision ^24^, and the proteomics workflows provided via the Galaxy platform ^25,26^) provide a graphical user interface where the user steps through the analysis manually; some additionally allow a summary report to be exported. Other software packages provide extensive collections of analysis functions, but require familiarity with programming to use (e.g. protti ^27^, POMA ^20^, SafeQuant ^28^, Proteus ^29^, DEP ^30^, prolfqua ^31^, MSstats ^32^). With einprot, we attempt to hit a middle ground by providing a collection of fully reproducible workflows that do not require extensive coding skills to run, yet return comprehensive, self-contained reports that contain all the executed code and are fully customizable if needed. In addition, by returning the data and results in a standardized format, the interoperability with other packages, including some of the ones mentioned above, is simplified and allows the user to take advantage of a vast ecosystem of tools. One such example is the direct interface to iSEE, which lets the user interactively explore all the results and processed values generated by and documented in the reproducible workflows. In this way, einprot is used productively by the proteomics and protein analysis facility at FMI and as the basis for published and submitted articles ^33^.

## AVAILABILITY AND INSTALLATION

einprot is available from GitHub (https://github.com/fmicompbio/einprot) and can be installed using the standard installation processes for R packages (e.g., using the remotes package ^34^). Documentation and example usage are available from https://fmicompbio.github.io/einprot/. A collection of example reports can be browsed at https://csoneson.github.io/einprot_examples/.

## ACKNOWLEDGMENTS

The authors would like to thank Merle Skribbe, Seraina Steiger, Thomas Welte, Patrick Matthias, Helge Grosshans and Marc Bühler for testing and feedback on the software, and Laurent Gatto for valuable discussions. This work was supported by the Novartis Research Foundation. The funding body did not have any role in the design of the study, the collection, analysis, and interpretation of data, or in writing the manuscript.

## Notes

### Competing Interest Statement

The authors have declared no competing interest.

## REFERENCES

[1] Hui Peng, He Wang, Weijia Kong, Jinyan Li, and Wilson Wen Bin Goh. Optimizing proteomics data differential expression analysis via High-Performing rules and ensemble inference. bioRxiv doi:10.1101/2023.06.26.546625, 2023.

[2] Jürgen Cox and Matthias Mann. MaxQuant enables high peptide identification rates, individualized p.p.b.-range mass accuracies and proteome-wide protein quantification. Nature Biotechnology, 26(12):1367–1372, 2008.

[3] Andy T Kong, Felipe V Leprevost, Dmitry M Avtonomov, Dattatreya Mellacheruvu, and Alexey I Nesvizhskii. MSFragger: ultrafast and comprehensive peptide identification in mass spectrometry-based proteomics. Nature Methods, 14(5):513–520, 2017.

[4] Benjamin C Orsburn. Proteome Discoverer-A community enhanced data processing suite for protein informatics. Proteomes, 9(1), 2021.

[5] Yihui Xie, J.J. Allaire, and Garrett Grolemund. R Markdown: The Definitive Guide. Chapman and Hall/CRC, Boca Raton, Florida, 2018.

[6] Xiu Nie, Liujia Qian, Rui Sun, Bo Huang, Xiaochuan Dong, Qi Xiao, Qiushi Zhang, Tian Lu, Liang Yue, Shuo Chen, Xiang Li, Yaoting Sun, Lu Li, Luang Xu, Yan Li, Ming Yang, Zhangzhi Xue, Shuang Liang, Xuan Ding, Chunhui Yuan, Li Peng, Wei Liu, Xiao Yi, Mengge Lyu, Guixiang Xiao, Xia Xu, Weigang Ge, Jiale He, Jun Fan, Junhua Wu, Meng Luo, Xiaona Chang, Huaxiong Pan, Xue Cai, Junjie Zhou, Jing Yu, Huanhuan Gao, Mingxing Xie, Sihua Wang, Guan Ruan, Hao Chen, Hua Su, Heng Mei, Danju Luo, Dashi Zhao, Fei Xu, Yan Li, Yi Zhu, Jiahong Xia, Yu Hu, and Tiannan Guo. Multi-organ proteomic landscape of COVID-19 autopsies. Cell, 184(3):775–791.e14, 2021.

[7] Tianen He, Youqi Liu, Yan Zhou, Lu Li, He Wang, Shanjun Chen, Jinlong Gao, Wenhao Jiang, Yi Yu, Weigang Ge, Hui-Yin Chang, Ziquan Fan, Alexey I Nesvizhskii, Tiannan Guo, and Yaoting Sun. Comparative evaluation of Proteome Discoverer and FragPipe for the TMT-Based proteome quantification. Journal of Proteome Research, 21(12):3007–3015, 2022.

[8] Veronika Ostapcuk, Fabio Mohn, Sarah H Carl, Anja Basters, Daniel Hess, Vytautas Iesmantavicius, Lisa Lampersberger, Matyas Flemr, Aparna Pandey, Nicolas H Thomä, Joerg Betschinger, and Marc Bühler. Activity-dependent neuroprotective protein recruits HP1 and CHD4 to control lineage-specifying genes. Nature, 557(7707):739–743, 2018.

[9] Michael Ashburner, Catherine A Ball, Judith A Blake, David Botstein, Heather Butler, J Michael Cherry, Allan P Davis, Kara Dolinski, Selina S Dwight, Janan T Eppig, Midori A Harris, David P Hill, Laurie Issel-Tarver, Andrew Kasarskis, Suzanna Lewis, John C Matese, Joel E Richardson, Martin Ringwald, Gerald M Rubin, and Gavin Sherlock. Gene ontology: tool for the unification of biology. Nature Genetics, 25(1):25–29, 2000.

[10] Gene Ontology Consortium, Suzi A Aleksander, James Balhoff, Seth Carbon, J Michael Cherry, Harold J Drabkin, Dustin Ebert, Marc Feuermann, Pascale Gaudet, Nomi L Harris, David P Hill, Raymond Lee, Huaiyu Mi, Sierra Moxon, Christopher J Mungall, Anushya Muruganugan, Tremayne Mushayahama, Paul W Sternberg, Paul D Thomas, Kimberly Van Auken, Jolene Ramsey, Deborah A Siegele, Rex L Chisholm, Petra Fey, Maria Cristina Aspromonte, Maria Victoria Nugnes, Federica Quaglia, Silvio Tosatto, Michelle Giglio, Suvarna Nadendla, Giulia Antonazzo, Helen Attrill, Gil Dos Santos, Steven Marygold, Victor Strelets, Christopher J Tabone, Jim Thurmond, Pinglei Zhou, Saadullah H Ahmed, Praoparn Asanitthong, Diana Luna Buitrago, Meltem N Erdol, Matthew C Gage, Mohamed Ali Kadhum, Kan Yan Chloe Li, Miao Long, Aleksandra Michalak, Angeline Pesala, Armalya Pritazahra, Shirin C C Saverimuttu, Renzhi Su, Kate E Thurlow, Ruth C Lovering, Colin Logie, Snezhana Oliferenko, Judith Blake, Karen Christie, Lori Corbani, Mary E Dolan, Harold J Drabkin, David P Hill, Li Ni, Dmitry Sitnikov, Cynthia Smith, Alayne Cuzick, James Seager, Laurel Cooper, Justin Elser, Pankaj Jaiswal, Parul Gupta, Pankaj Jaiswal, Sushma Naithani, Manuel Lera-Ramirez, Kim Rutherford, Valerie Wood, Jeffrey L De Pons, Melinda R Dwinell, G Thomas Hayman, Mary L Kaldunski, Anne E Kwitek, Stanley J F Laulederkind, Marek A Tutaj, Mahima Vedi, Shur-Jen Wang, Peter D’Eustachio, Lucila Aimo, Kristian Axelsen, Alan Bridge, Nevila Hyka-Nouspikel, Anne Morgat, Suzi A Aleksander, J Michael Cherry, Stacia R Engel, Kalpana Karra, Stuart R Miyasato, Robert S Nash, Marek S Skrzypek, Shuai Weng, Edith D Wong, Erika Bakker, Tanya Z Berardini, Leonore Reiser, Andrea Auchincloss, Kristian Axelsen, Ghislaine Argoud-Puy, Marie-Claude Blatter, Emmanuel Boutet, Lionel Breuza, Alan Bridge, Cristina Casals-Casas, Elisabeth Coudert, Anne Estreicher, Maria Livia Famiglietti, Marc Feuermann, Arnaud Gos, Nadine Gruaz-Gumowski, Chantal Hulo, Nevila Hyka-Nouspikel, Florence Jungo, Philippe Le Mercier, Damien Lieberherr, Patrick Masson, Anne Morgat, Ivo Pedruzzi, Lucille Pourcel, Sylvain Poux, Catherine Rivoire, Shyamala Sundaram, Alex Bateman, Emily Bowler-Barnett, Hema Bye-A-Jee, Paul Denny, Alexandr Ignatchenko, Rizwan Ishtiaq, Antonia Lock, Yvonne Lussi, Michele Magrane, Maria J Martin, Sandra Orchard, Pedro Raposo, Elena Speretta, Nidhi Tyagi, Kate Warner, Rossana Zaru, Alexander D Diehl, Raymond Lee, Juancarlos Chan, Stavros Diamantakis, Daniela Raciti, Magdalena Zarowiecki, Malcolm Fisher, Christina James-Zorn, Virgilio Ponferrada, Aaron Zorn, Sridhar Ramachandran, Leyla Ruzicka, and Monte Westerfield. The gene ontology knowledgebase in 2023. Genetics, 224(1), 2023.

[11] Madalina Giurgiu, Julian Reinhard, Barbara Brauner, Irmtraud Dunger-Kaltenbach, Gisela Fobo, Goar Frishman, Corinna Montrone, and Andreas Ruepp. CORUM: the comprehensive resource of mammalian protein complexes-2019. Nucleic Acids Research, 47(D1):D559–D563, 2019.

[12] Midori A Harris, Kim M Rutherford, Jacqueline Hayles, Antonia Lock, Jürg Bähler, Stephen G Oliver, Juan Mata, and Valerie Wood. Fission stories: using PomBase to understand Schizosaccharomyces pombe biology. Genetics, 220(4), 2022.

[13] Shuye Pu, Jessica Wong, Brian Turner, Emerson Cho, and Shoshana J Wodak. Up-to-date catalogues of yeast protein complexes. Nucleic Acids Research, 37(3):825–831, 2009.

[14] Igor Dolgalev. babelgene: Gene Orthologs for Model Organisms in a Tidy Data Format, 2022. R package version 22.9.

[15] Wolfgang Huber, Vincent J Carey, Robert Gentleman, Simon Anders, Marc Carlson, Benilton S Carvalho, Hector Corrada Bravo, Sean Davis, Laurent Gatto, Thomas Girke, Raphael Gottardo, Florian Hahne, Kasper D Hansen, Rafael A Irizarry, Michael Lawrence, Michael I Love, James MacDonald, Valerie Obenchain, Andrzej K Oleś, Hervé Pagès, Alejandro Reyes, Paul Shannon, Gordon K Smyth, Dan Tenenbaum, Levi Waldron, and Martin Morgan. Orchestrating high-throughput genomic analysis with Bioconductor. Nature Methods, 12(2):115–121, 2015.

[16] Robert A Amezquita, Aaron T L Lun, Etienne Becht, Vince J Carey, Lindsay N Carpp, Ludwig Geistlinger, Federico Marini, Kevin Rue-Albrecht, Davide Risso, Charlotte Soneson, Levi Waldron, Hervé Pagès, Mike L Smith, Wolfgang Huber, Martin Morgan, Raphael Gottardo, and Stephanie C Hicks. Orchestrating single-cell analysis with Bioconductor. Nature Methods, 17(2):137–145, 2020.

[17] Kevin Rue-Albrecht, Federico Marini, Charlotte Soneson, and Aaron T L Lun. iSEE: Interactive SummarizedExperiment Explorer. F1000 Research, 7:741, 2018.

[18] Mingze Bai, Jingwen Deng, Chengxin Dai, Julianus Pfeuffer, Timo Sachsenberg, and Yasset Perez-Riverol. LFQ-Based peptide and protein intensity differential expression analysis. Journal of Proteome Research, 22(6):2114–2123, 2023.

[19] Stefka Tyanova, Tikira Temu, Pavel Sinitcyn, Arthur Carlson, Marco Y Hein, Tamar Geiger, Matthias Mann, and Jürgen Cox. The Perseus computational platform for comprehensive analysis of (prote)omics data. Nature Methods, 13(9):731–740, 2016.

[20] Pol Castellano-Escuder, Raúl González-Domínguez, Francesc Carmona-Pontaque, Cristina Andrés-Lacueva, and Alex Sánchez-Pla. POMAShiny: A user-friendly web-based workflow for metabolomics and proteomics data analysis. PLoS Computational Biology, 17(7):e1009148, 2021.

[21] Anup D Shah, Robert J A Goode, Cheng Huang, David R Powell, and Ralf B Schittenhelm. LFQ-Analyst: An Easy-To-Use interactive web platform to analyze and visualize Label-Free proteomics data preprocessed with MaxQuant. Journal of Proteome Research, 19(1):204–211, 2020.

[22] Sebastian Didusch, Moritz Madern, Markus Hartl, and Manuela Baccarini. amica: an interactive and user-friendly web-platform for the analysis of proteomics data. BMC Genomics, 23(1):817, 2022.

[23] Milena Kraus, Mariet Mathew Stephen, and Matthieu-P Schapranow. Eatomics: Shiny exploration of quantitative proteomics data. Journal of Proteome Research, 20(1):1070–1078, 2021.

[24] James Luke Gallant, Tiaan Heunis, Samantha Leigh Sampson, and Wilbert Bitter. ProVision: a web-based platform for rapid analysis of proteomics data processed by MaxQuant. Bioinformatics, 36(19):4965–4967, 2020.

[25] Galaxy Community. The Galaxy platform for accessible, reproducible and collaborative biomedical analyses: 2022 update. Nucleic Acids Research, 50(W1):W345–W351, 2022.

[26] Saskia Hiltemann, Helena Rasche, Simon Gladman, Hans-Rudolf Hotz, Delphine Larivière, Daniel Blankenberg, Pratik D Jagtap, Thomas Wollmann, Anthony Bretaudeau, Nadia Goué, Timothy J Griffin, Coline Royaux, Yvan Le Bras, Subina Mehta, Anna Syme, Frederik Coppens, Bert Droesbeke, Nicola Soranzo, Wendi Bacon, Fotis Psomopoulos, Cristóbal Gallardo-Alba, John Davis, Melanie Christine Föll, Matthias Fahrner, Maria A Doyle, Beatriz Serrano-Solano, Anne Claire Fouilloux, Peter van Heusden, Wolfgang Maier, Dave Clements, Florian Heyl, Galaxy Training Network, Björn Grüning, and Bérénice Batut. Galaxy training: A powerful framework for teaching! PLoS Computational Biology, 19(1):e1010752, 2023.

[27] Jan-Philipp Quast, Dina Schuster, and Paola Picotti. protti: an R package for comprehensive data analysis of peptide- and protein-centric bottom-up proteomics data. Bioinformatics Advances, 2(1):vbab041, 2021.

[28] Timo Glatter, Christina Ludwig, Erik Ahrné, Ruedi Aebersold, Albert J R Heck, and Alexander Schmidt. Large-scale quantitative assessment of different in-solution protein digestion protocols reveals superior cleavage efficiency of tandem Lys-C/trypsin proteolysis over trypsin digestion. Journal of Proteome Research, 11(11):5145–5156, 2012.

[29] Marek Gierlinski, Francesco Gastaldello, Chris Cole, and Geoffrey J Barton. Proteus: an R package for downstream analysis of MaxQuant output. bioRxiv doi:10.1101/416511, 2018.

[30] Xiaofei Zhang, Arne H Smits, Gabrielle B A van Tilburg, Huib Ovaa, Wolfgang Huber, and Michiel Vermeulen. Proteome-wide identification of ubiquitin interactions using UbIA-MS. Nature Protocols, 13(3):530–550, 2018.

[31] Witold E Wolski, Paolo Nanni, Jonas Grossmann, Maria d’Errico, Ralph Schlapbach, and Christian Panse. prolfqua: A comprehensive R-Package for proteomics differential expression analysis. Journal of Proteome Research, 22(4):1092–1104, 2023.

[32] Meena Choi, Ching-Yun Chang, Timothy Clough, Daniel Broudy, Trevor Killeen, Brendan MacLean, and Olga Vitek. MSstats: an R package for statistical analysis of quantitative mass spectrometry-based proteomic experiments. Bioinformatics, 30(17):2524–2526, 2014.

[33] Thomas Welte, Alison Goulois, Michael B Stadler, Daniel Hess, Charlotte Soneson, Anca Neagu, Chiara Azzi, Marlena J Wisser, Jan Seebacher, Isabel Schmidt, David Estoppey, Florian Nigsch, John Reece-Hoyes, Dominic Hoepfner, and Helge Großhans. Convergence of multiple RNA-silencing pathways on GW182/TNRC6. Molecular Cell, 2023.

[34] Gábor Csárdi, Jim Hester, Hadley Wickham, Winston Chang, Martin Morgan, and Dan Tenenbaum. remotes: R Package Installation from Remote Repositories, Including ‘GitHub’, 2021. R package version 2.4.2.

